# Effects of Polyunsaturated Fatty Acids and Selenoneine on Growth and Lipidomic Profiles in Patient-Derived Prostate Cancer Organoids

**DOI:** 10.64898/2026.09.04.749239

**Authors:** Mame Sokhna Sylla, Cynthia Jobin, Line Berthiaume, Lucie Leclair, Gabriel Lachance, Karine Robitaille, Claire Ménard, Chantal Atallah, Frédéric Pouliot, Pierre Ayotte, Yves Fradet, Vincent Fradet, Étienne Audet-Walsh

**Author notes:** Corresponding Authors: Étienne Audet-Walsh, Centre de recherche du CHU de Québec, 2705 Boulevard Laurier, room R-4714, Québec City, QC, Canada, G1V 4G2., Vincent Fradet, Centre de recherche du CHU de Québec-Université Laval, 2250, boulevard Henri Bourassa, room G4.610, Québec City, QC, Canada, G1J 1Z4., Yves Fradet, Centre de recherche du CHU de Québec-Université Laval, 2250, boulevard Henri Bourassa, room G4.610, Québec City, QC, Canada, G1J 1Z4.

## Abstract

Prostate cancer (PCa) is the second most frequent cancer in men and represents a major public health issue worldwide. To complement current therapeutic approaches, our work aims to understand the impact of nutritional interventions on PCa evolution. Recent studies showed that supplementation with omega-3 fatty acids reduces tumour growth in PCa mouse models. Several beneficial effects were observed, particularly on the gut microbiome, suggesting indirect effects on tumours, and whether these fatty acids have a direct molecular impact on the prostate remains elusive. Here, we hypothesized that the prostate directly takes up dietary fatty acids to reprogram metabolism, which could contribute to the beneficial effects on cancer treatment. To test this hypothesis, five series of patient-derived organoids (PDOs), with paired lines originating from normal and tumour samples, were treated with omega-3 and omega-6 fatty acids. While heterogeneity was observed in growth capacities of PDO lines, omega-3 supplementation overall decreased PDO growth, with a stronger decrease in PCa lines. This effect was stronger when co-treated with selenoneine, a nutrient enriched in some marine foods with high omega-3 content. Lipidomic experiments were then performed, revealing that PDOs can take up fatty acids, both omega-3 and omega-6. Furthermore, PDOs could metabolize omega-3 fatty acids into downstream metabolites, even if high inter-patient heterogeneity was observed. Altogether, this study, using PDOs as a working model to study the human prostate, provides a better understanding of the molecular impacts of fatty acid supplementation on prostate and PCa biology for the development of future nutritional interventions.

## Introduction

Prostate cancer (PCa) is the second leading cause of cancer mortality in men worldwide^1^, highlighting this disease as a major public health issue. In this context, discovering new treatments that either help prevent the disease, improve patients’ quality of life, or prolong patients’ life expectancy is a critical clinical need. While Caucasians and African Americans frequently develop PCa, it was shown that Asians are less likely to develop PCa^2,3^. While this difference might have a genetic component, it is also clear now that there is an environmental component as well. Indeed, Japanese men in Japan have a lower PCa incidence than citizens of the United States of America, whereas those who migrated to North America harbour a much higher PCa incidence, indicating that changes in their environment, including their nutrition, significantly affect PCa development^3,4^. In Canada, PCa accounts for approximately 23% of all diagnosed cancers in men^5^. Interestingly, Canadian Inuit have a 10- to 5-fold lower risk of developing PCa relative to Caucasians or non-Inuit people^6,7^. Histological analyses from 27 autopsies showed no latent carcinoma in older individuals^7^. The diet of the Canadian Inuit is generally characterized by a high daily intake of omega-3 polyunsaturated fatty acids (PUFAs) and a lower omega-6/omega-3 ratio ^8,9^. In Nunavik, the Canadian Inuit diet is also highly enriched in selenoneine, which is abundant in some traditional marine foods^10^. Given that Inuit have elevated blood levels of selenoneine^11,12^ and a low incidence of PCa, dietary selenoneine intake may reduce PCa incidence, as shown for colorectal cancer^10^. Therefore, an omega-3-enriched diet supplemented with selenoneine could explain, at least in part, the lower risk of PCa incidence among Inuit populations.

In the daily diet, a variety of fatty acids are consumed, including omega-3, omega-6, omega-9, and saturated fatty acids, which serve distinct functions in cellular biology. Omega-3 and omega-6 are essential PUFAs that humans cannot produce. The primary sources of omega-3 fatty acids are marine foods, such as fish and sea mammals^13,14^, where eicosapentaenoic acid (EPA, 20:5 ω-3) and docosahexaenoic acid (DHA, 22:6 ω-3) are abundant^14^. Omega-3 fatty acids are derived from alpha-linolenic acid (ALA, 18:3 ω-3), mainly found in plants and some vegetable oils. ALA is desaturated to stearidonic acid (SDA, 18:4 ω-3) before being elongated into EPA, docosapentaenoic acid (DPA, 22:5 ω-3) and DHA^13^. Omega-6 fatty acids, such as linoleic acid (LA, 18:2 ω-6), are a family of PUFAs abundant in vegetable oils. LA can be desaturated into gamma-linolenic acid (GLA, 18:3 ω-6). Encoded by the fatty acid elongases *ELOVL5* and *ELOVL2*, GLA is elongated into dihomo-gamma-linolenic acid (DGLA, 20:3 ω-6), which can be desaturated into arachidonic acid (AA, 20:4 ω-6). Subsequently, AA is elongated into adrenic acid (22:4 ω-6)^15,16^. These PUFAs are generally free fatty acids, ethyl esters and mono-acylglycerol (MAG), the latter being taken up more easily by cells^17^.

Omega-3 fatty acids generally have anti-inflammatory effects and promote apoptosis in cancer cells^18–20^. In that context, omega-3 fatty acids are thought to have tumour suppressor functions. Indeed, while MAG-EPA reduced PCa tumour volumes in xenografted or transgenic mice, patients were downgraded, highlighting lower progression of PCa^21,22^. Conversely, omega-6 PUFAs can be metabolized into oxylipins that promote oncogenic pathways in PCa^23^. These results suggest that an omega-6-enriched diet has oncogenic functions. Consequently, a low-fat diet might reduce PCa progression by decreasing the omega-6/omega-3 ratio^24^. Taken together, an omega-3-enriched diet might protect against PCa, but the functional effects of omega-3 and omega-6 PUFAs on lipid metabolism in the prostate and PCa remain unclear.

The current study aimed to clarify the roles of omega-3 fatty acids in the prostate and in tumour cell proliferation. To achieve this, we used patient-derived organoids (PDOs) from paired normal and tumour lines from the same patient, which were treated with different types of fatty acids. Despite patient heterogeneity, omega-3 significantly reduces cell viability in both normal and tumour conditions. Lipidomic analyses revealed that PDOs can take up omega-3 fatty acids and metabolize them into new fatty acids specific to either normal or tumour states, likely rewiring lipid metabolism to support PCa cell proliferation. Thus, while omega-3 and omega-6 fatty acids might have indirect effects on PCa progression, such as modulating the gut microbiome, they could also have direct impacts on the prostate epithelium.

## Results

### Supplementation with omega-3 PUFAs to PDOs from primary PCa patients reduces PCa cell growth

The clinical characteristics of the five patients included in the current study are shown in Fig. 1A. All patients underwent radical prostatectomy. At surgery, fresh tumour samples were isolated and processed in primary culture to generate five series of PDOs (Fig. 1B) using a previously described protocol^25^. For each series, normal and PCa PDO lines were cultured in parallel, being isolated from normal vs tumour regions under the supervision of a pathologist. The average prostate weight was 46 g. Four patients had tumours with a Gleason score of 7, and one had a Gleason score of 6. Two patients were classified as pathological stage T2, while three were T3a. No patients had lymph node invasion. As expected, heterogeneity was observed between patients and between normal and tumour PDO lines (Fig. 1C-D). Compared with normal conditions, three PCa lines grew faster and increased in size after 14 days in culture.

**Figure 1.**
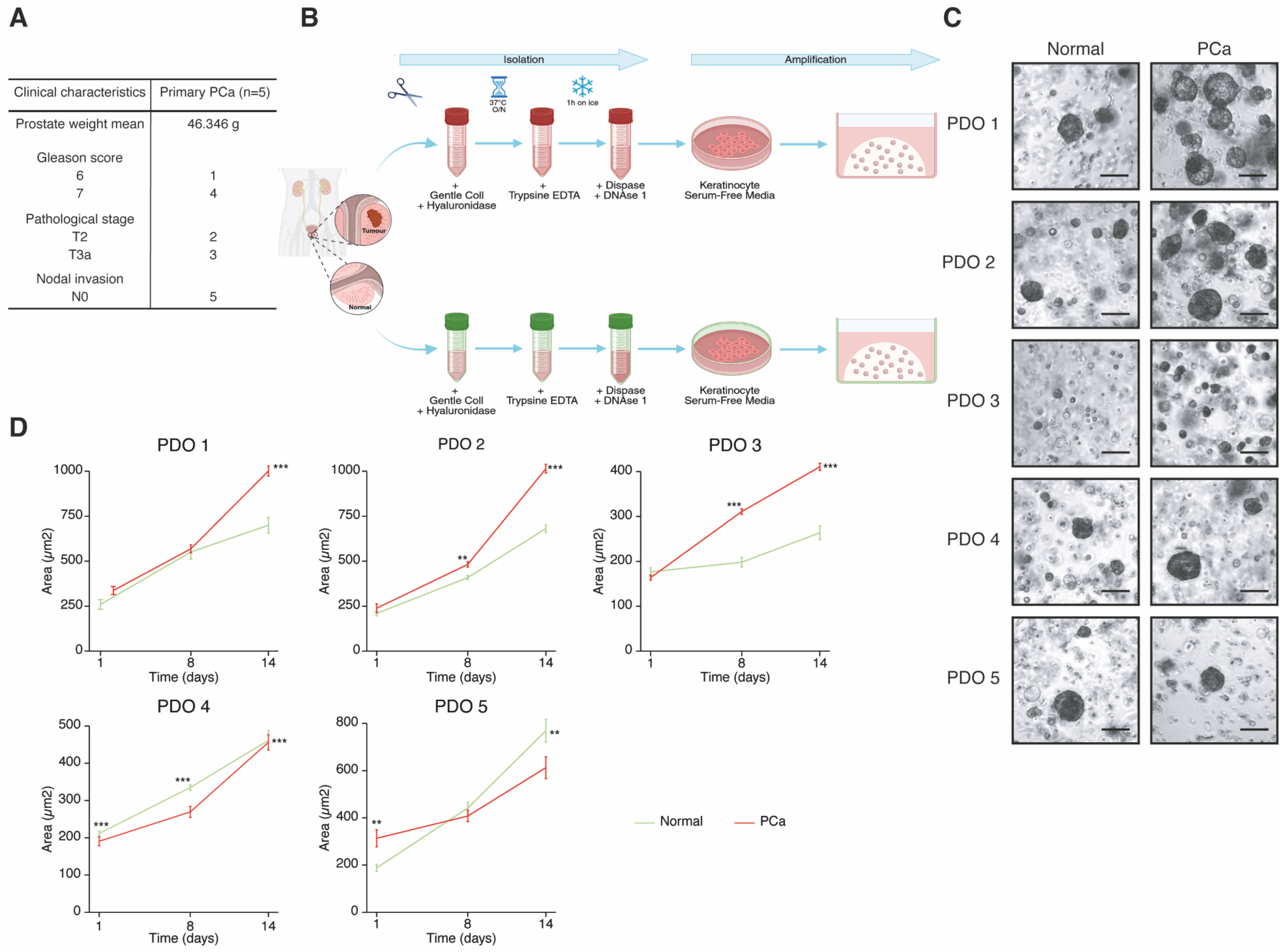
PDOs from primary PCa patients recapitulate patient heterogeneity. **(A)** Clinical characteristics of patients who underwent surgery and for whom prostate samples were received for primary culture. **(B)** The experimental design to generate PDO lines: Normal and PCa adjacent samples were isolated using different digestive enzymes before being amplified in 2D and 3D, respectively. **(C)** PDOs brightfield images taken at D14 in both normal and PCa conditions. **(D)** PDO quantification between normal and tumour conditions at D14. Treatments were normalized to the mean of the controls of each PDO. *P < 0.05, **P < 0.01, and ***P < 0.001, by wilcoxon-test adjusted with Bonferroni.

Using these five PDO series, the impact of omega-3 and omega-6 fatty acid supplementation was then tested (following the experimental design shown in Fig. 1B). In brief, 24h after seeding in Matrigel drops, PDOs were treated with media supplemented with MAG-EPA, with and without selenoneine, or with MAG-AA. PDOs were then cultured until they reached their maximal size at 14 days, at which point cell viability was measured (Fig. 2A-C). Again, inter-patient heterogeneity was observed in PDO growth, as visualized using brightfield microscopy (Fig. 2A). In normal prostate PDOs, MAG-EPA significantly decreased cell viability in PDO lines 4 and 5 (Fig. 2B). A trend for lower viability was also observed in other PDO lines, and, when all five lines were combined, MAG-EPA was associated with a significantly lower level of cell viability compared to controls (Fig. 2C).

**Figure 2.**
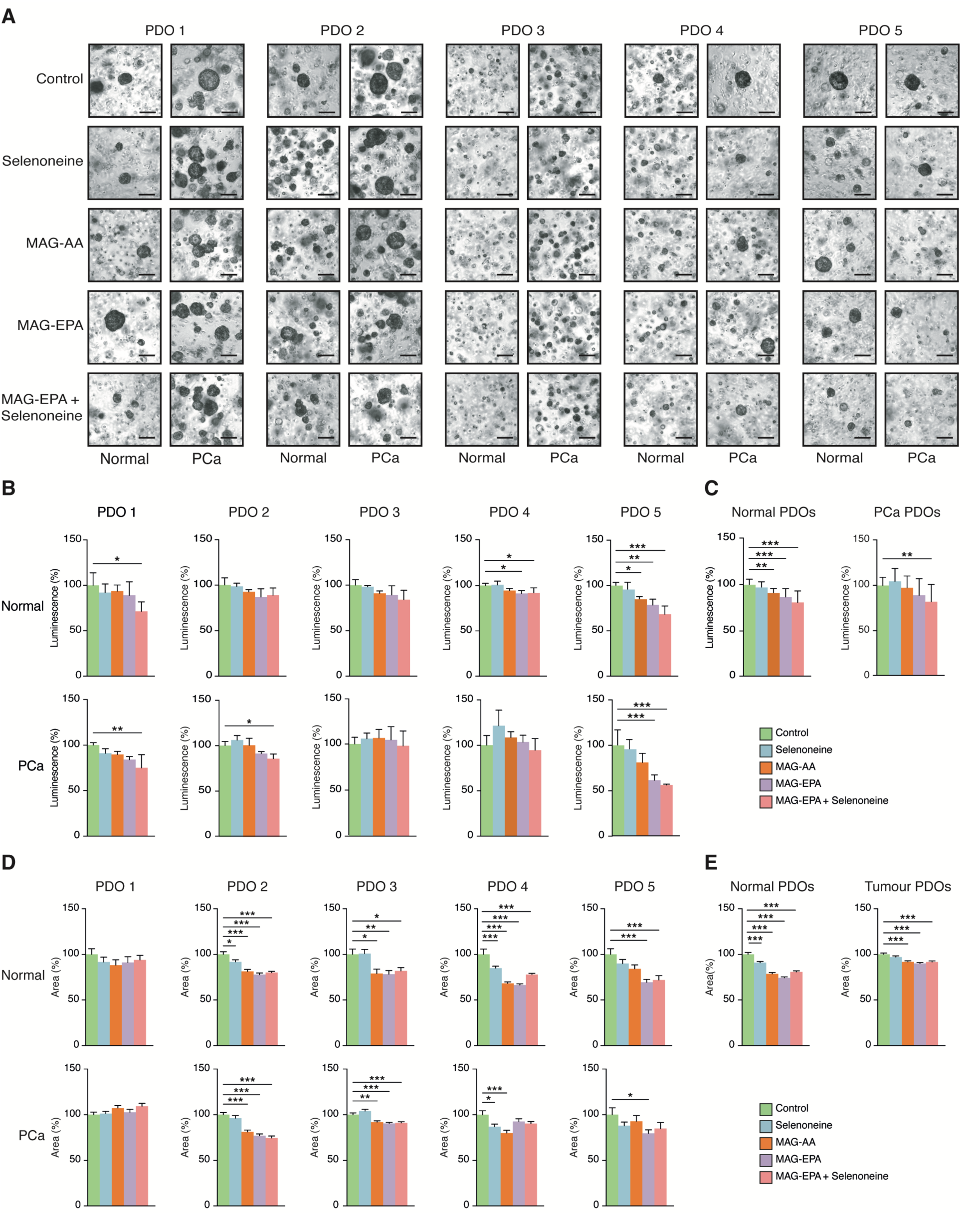
Supplementation with omega-3 PUFAs to PDOs reduces PCa cell growth and proliferation. **(A)** Omega-3 PUFAs modulate PDO growth compared to other treatments and controls. Brightfield images were taken at D14 for each treatment. **(B-C)** MAG-EPA reduces the viability of PDO lines. **(B)** The effects of treatments on the viability of each PDO. **(C)** The effects of treatments on the viability of combined PDOs in normal and PCa conditions. Data are shown as mean + SD (n = 3 replicates per condition). **(D-E)** Omega-3 fatty acids induce cell proliferation in **(D)** each PDO and **(E)** after merging all PDO lines. Data were normalized to the controls’ means. Data are shown as mean + SEM (n = 2 images per droplet per treatment per condition for a total of 2 droplets for each PDO). *P < 0.05, **P < 0.01, and ***P < 0.001, by one-way ANOVA.

For PCa PDOs, in individual lines, only MAG-EPA + selenoneine significantly decreased cell viability, being significant in the individual lines 1, 2, and 5 (Fig. 2B). As for normal prostate PDOs, combining all the lines showed a significant decrease in cell viability solely when MAG-EPA was combined with selenoneine (Fig. 2C). While no other treatments were significantly different compared to controls in PCa PDOs, the combination of all 5 lines together also revealed a significant decrease in cell viability following treatment with MAG-AA and MAG-EPA in normal prostate PDOs, suggesting a higher sensitivity of normal prostate epithelial cells to fatty acids (Fig. 2C). These results also suggest that tumour PDOs are more tolerant to omega-6 treatment.

Finally, organoid size was quantified after 14 days in culture to assess the effects of the treatments on organoid growth (Fig. 2D). For 4 normal PDO lines, supplementation with omega-6 and omega-3 fatty acids significantly decreased organoid growth. MAG-EPA, supplemented or not with selenoneine, significantly reduced the size of 3 series of PCa PDOs. When all PDOs were combined, either MAG-AA or MAG-EPA significantly decreased PDO size (Fig. 2E).

### Omega-3 supplementation modulates the lipid composition of PDOs

Next, lipidomic analyses were performed on PDO series 1, 2, and 4, for which enough organoid biological material was available. In PDO line 1, lipidomics revealed changes in the lipid profile in both normal and tumour conditions. In normal samples, while organoids treated with selenoneine clustered with controls, those treated with MAG-AA, MAG-EPA, or MAG-EPA + selenoneine clustered apart from the controls, indicating that the treatment altered their lipidome (Fig. 3A). A similar pattern was also observed for the PCa PDO line 1 (Fig. 3B), again highlighting a different lipidomic profile. In both conditions, MAG-EPA increased the levels of specific omega-3 fatty acids, EPA (20:5 ω-3), as expected, DPA (22:5 ω-3), DHA (22:6 ω-3), and ALA (18:3 ω-3) (Fig. 3C-D). Some fatty acids, especially saturated ones such as myristic acid (14:0) and palmitic acid (16:0), were uniquely enriched in controls or samples supplemented with selenoneine. Unlike normal organoids, tumour PDO line 1 showed a decrease in the levels of specific omega-9 fatty acids, including gondoic acid (20:1 ω-9), erucic acid (22:1 ω-9), and nervonic acid (24:1 ω-9) following MAG-EPA supplementation (Fig. 3D). Additionally, omega-6 fatty acids such as GLA, LA, and cis-13 octadecenoic acid (18:1 ω-5), were also increased after MAG-EPA treatment. Interestingly, cis-8,11,14,17 eicosatetraenoic acid (ETA, 20:4 ω-3) was specifically increased in this PCa PDO line. Consequently, treatment of PDO line 1 with MAG-EPA (containing also low levels of ALA and DHA) induced lipidomic changes, indicating active uptake by prostate epithelial cells. In both normal and PCa PDOs, there was a significant increase in the omega-3/omega-6 ratio (Fig. 3E-F). The changes observed following treatment with MAG-AA were more subtle, increasing the levels of AA itself and 22:4 ω-6. These results suggest that omega-3 fatty acids, and to a lesser extent omega-6 fatty acids, modulated the lipid profiles in the context of PCa.

**Figure 3.**
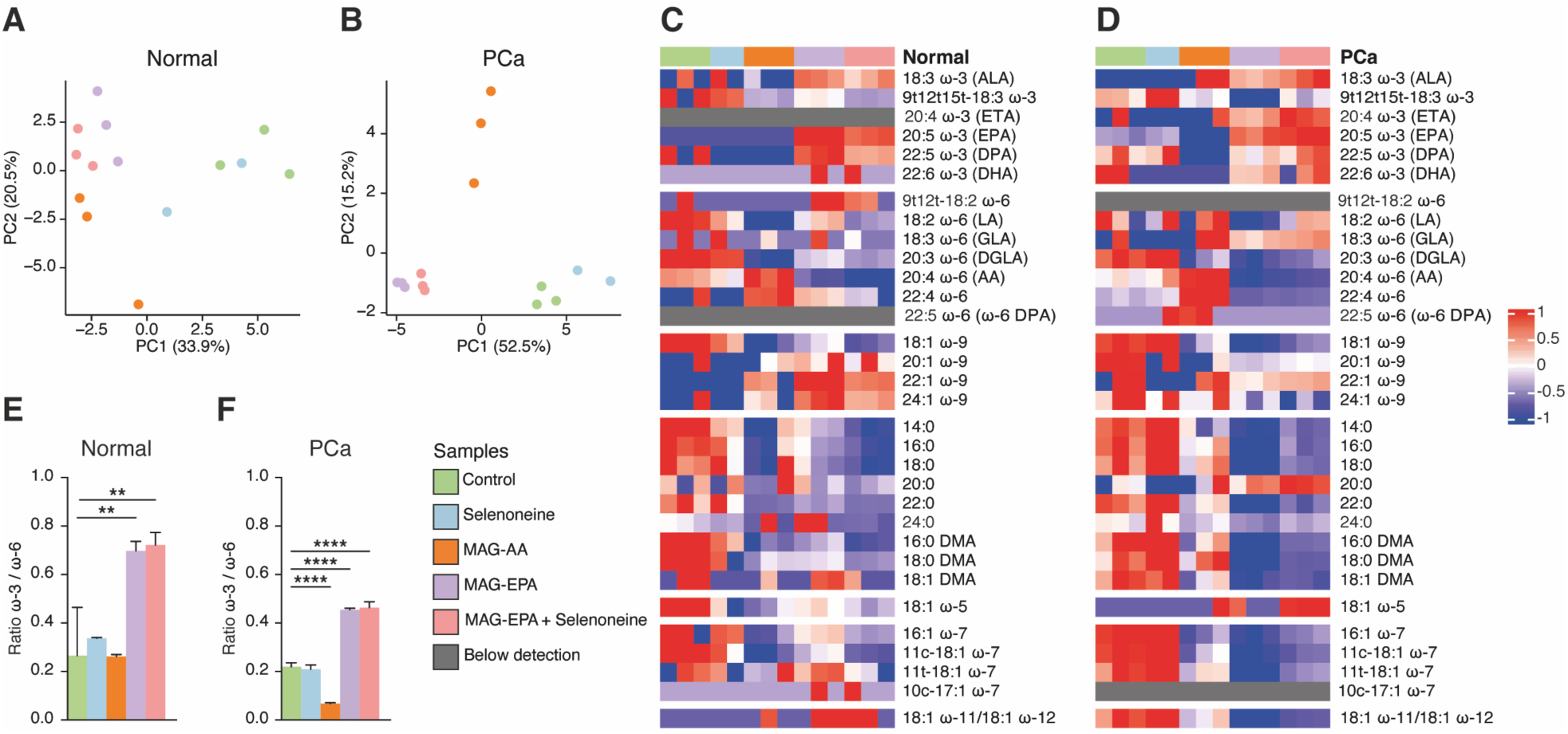
Omega-3 modulates the lipidome profile of PDO 1. **(A-B)** Principal Component Analyses (PCA) in the **(A)** normal and **(B)** PCa PDO line 1. **(C-D)** Heatmap of lipid composition following different treatments. **(C)** Normal PDO line. **(D)** PCa PDO line. **(E-F)** The effects of supplementation with fatty acids on the ratio omega-3/omega-6. **(E)** Normal PDO 1. **(F)** PCa PDO 1. N = 3 replicates per treatment per condition, except for selenoneine, with 2 replicates. *P < 0.05, **P < 0.01, and ***P < 0.001, by one-way ANOVA. Data are shown as ratio + SD.

The lipidomic profile of PDO line 2 was then studied in normal and PCa PDOs (Fig. 4A-B). In the normal PDO line 2, while treatment conditions tended to cluster together, controls clustered closer to the other treatment groups, suggesting a smaller effect on the lipidome (Fig. 4A). Of note, only organoids treated with MAG-AA clearly clustered apart. Treatment with MAG-EPA, similarly for the PDO line 1, mostly increased the levels of EPA, DPA, and ALA. The co-treatment of MAG-EPA with selenoneine had a major impact, and most fatty acids were increased further (Fig. 4C). Alongside omega-3 fatty acids, omega-6 fatty acids such as LA, DGLA, and GLA were also at higher levels in these samples. Omega-9 and saturated fatty acids were additionally found to be uniquely increased in omega-3 samples supplemented with selenoneine. In PCA analysis of the PCa line, organoids treated with MAG-EPA had the most well-defined cluster apart from the controls. In this line, while MAG-EPA led to a similar increase in EPA, DPA, and ALA, co-treatment with selenoneine decreased most fatty acids, rather than increasing them (Fig. 4D). As seen for the PDO line 1, MAG-AA treatment had more limited effects, increasing the levels of AA itself, along with 22:4 ω-6. In both normal and tumour conditions, the omega-3/omega-6 ratio was significantly increased following supplementation with MAG-EPA, while it decreased with MAG-AA (Fig. 4E-F).

**Figure 4.**
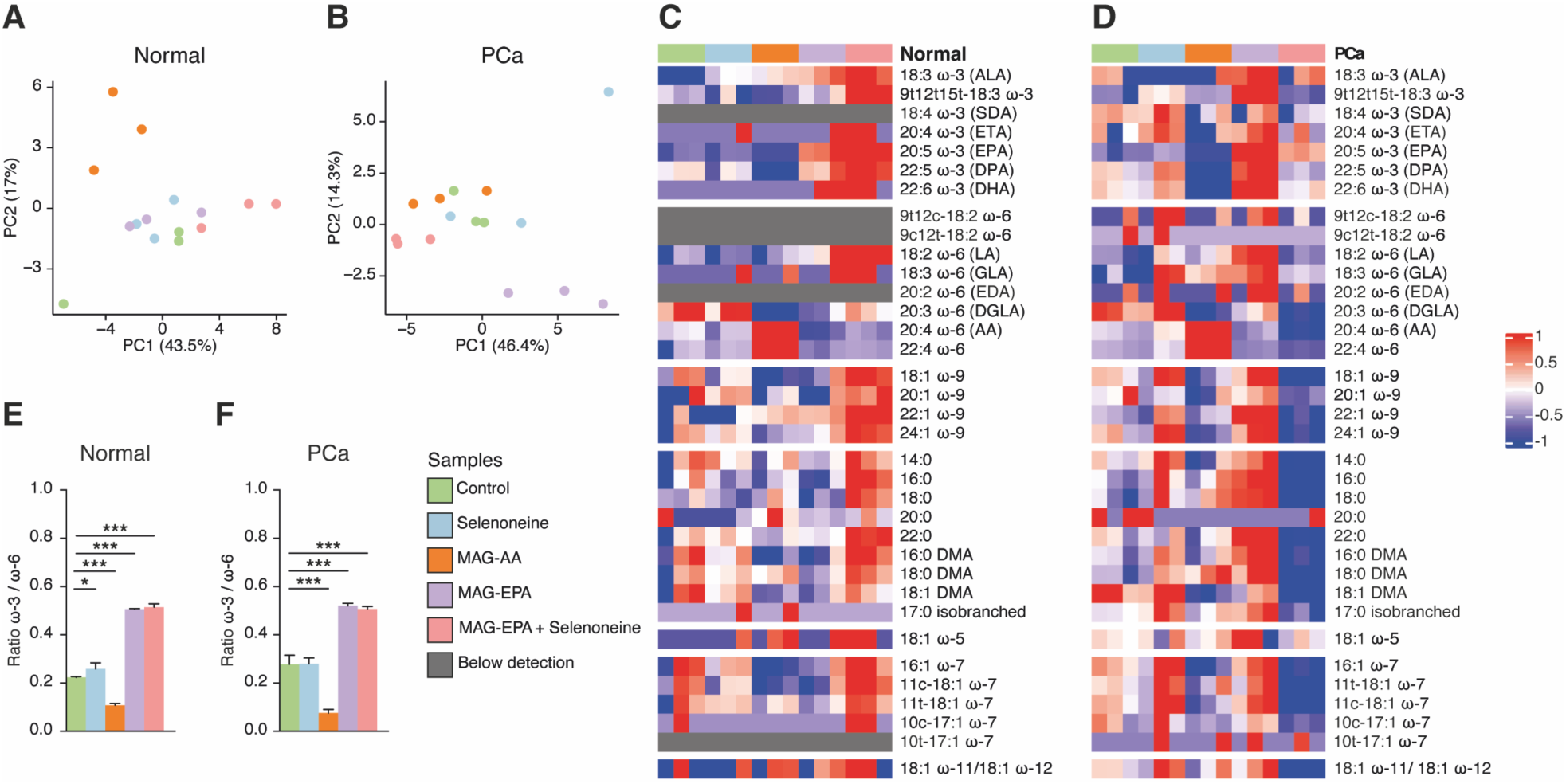
Supplementation with PUFAs modulates the lipid composition of PDO 2. **(A-B)** Principal Component Analyses in **(A)** normal and **(B)** PCa conditions. **(C-D)** Heatmap of lipid composition following different treatments. **(C)** Normal PDO line. **(D)** PCa PDO line. **(E-F)** The effects of supplementation with fatty acids on the ratio omega-3/omega-6. **(E)** Normal PDO 2. **(F)** PCa PDO 2. N = 3 replicates per treatment per condition. *P < 0.05, **P < 0.01, and ***P < 0.001, by one-way ANOVA. Data are shown as ratio + SD.

In the PDO line 4, the controls were best separated in the PCa line, suggesting that fatty acid treatments had a stronger effect, particularly in the tumour model (Fig. 5A-B). In the normal line 4 (Fig. 5C), as observed in previous lines (Fig. 3-4), supplementation with MAG-EPA increased the levels of EPA, DHA, and DPA. Similarly, these fatty acids were also highly increased in the PCa line (Fig. 5D). While gondoic acid (20:1 ω-9) was abundant in normal samples, it was absent in the PCa PDO line. However, MAG-AA induces enrichment of various fatty acid types, such as saturated fatty acids, in normal samples (Fig. 5C). While supplementing MAG-EPA with selenoneine did not significantly modulate the omega-3/omega-6 ratio, MAG-AA significantly decreased it (Fig. 5E-F). Thus, MAG-EPA, as seen with other PDO lines, mostly modified other omega-3 fatty acids, while distinct results were obtained for other fatty acids between the normal and tumour lines.

**Figure 5.**
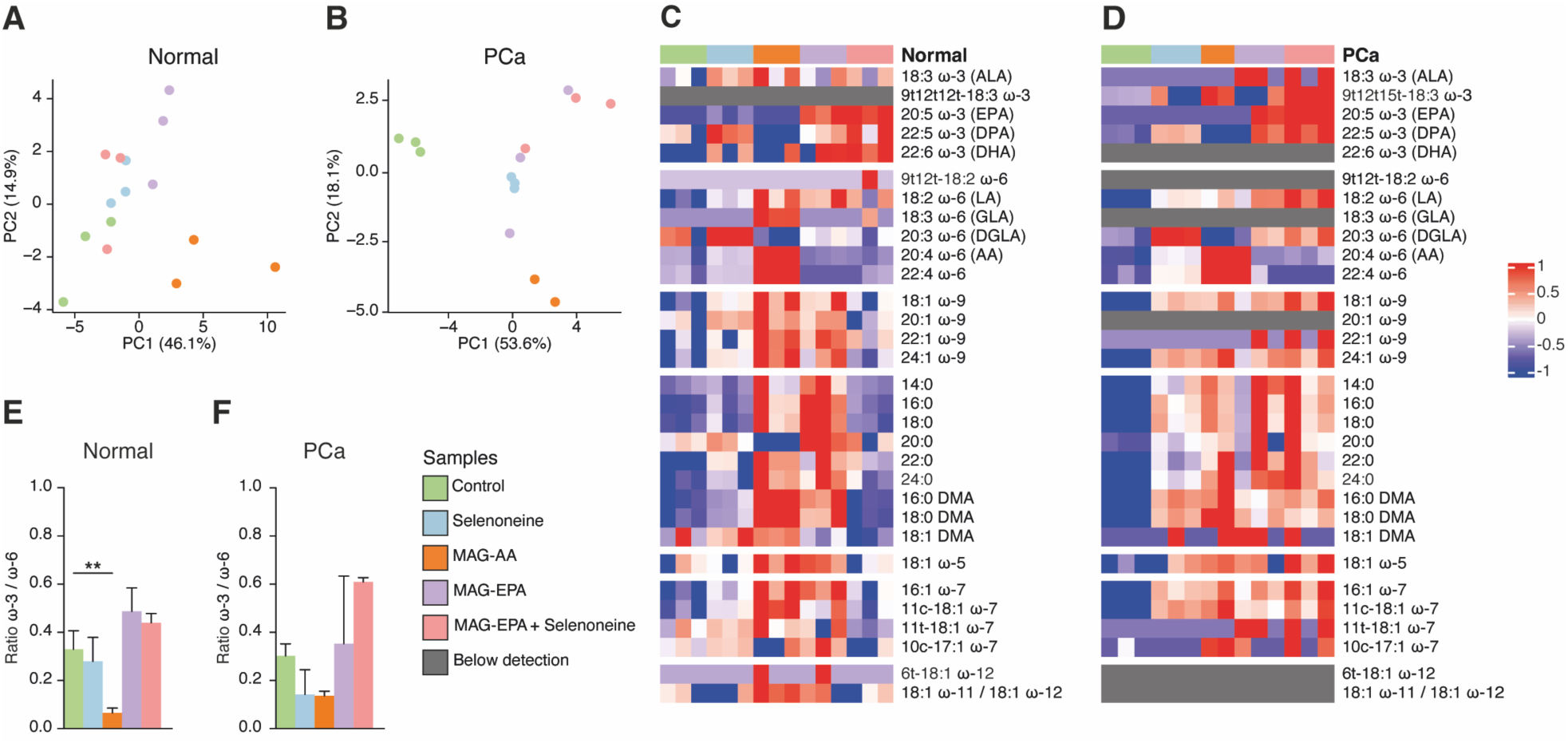
Treating with omega-3 PUFAs modulates the lipidome profile of PDO 4. **(A-B)** Principal Component Analyses in **(A)** normal and **(B)** PCa conditions. **(C-D)** Heatmap of lipid composition following different treatments. **(C)** Normal PDO line. **(D)** PCa PDO line. **(E-F)** The effects of supplementation with fatty acids on the ratio omega-3/omega-6. **(E)** Normal PDO 4. **(F)** PCa PDO 4. N = 3 replicates per treatment per condition, except for MAG-AA in the PCa condition, with 2 replicates. *P < 0.05, **P < 0.01, and ***P < 0.001, by one-way ANOVA. Data are shown as ratio + SD.

## Discussion

This study used PDOs to investigate the effects of fatty acids on prostate and PCa epithelial cells grown as three-dimensional organoid models. First, it was shown that MAG-EPA significantly reduced PDO viability, especially when combined with selenoneine. Moreover, omega-3 fatty acids significantly lowered PCa PDO growth. Lipidomic analyses indicated that MAG-EPA altered the lipid profile of PDOs, notably increasing levels of various omega-3 PUFAs and the omega-3/omega-6 ratio. Treatment with MAG-AA had a limited impact on the lipid profiles of PDOs, indicating some uptake but limited downstream metabolism.

Here, supplementation with MAG-EPA, notably combined with selenoneine, was shown to be more effective at inhibiting PCa PDO viability. These results align with previous studies using cell lines, which showed that treatment with omega-3 PUFAs, such as EPA, promotes apoptosis in cancer cells^26,27^. Cellular viability assays in DU145, an AR-negative cell line, demonstrated that omega-3 fatty acids, specifically EPA and DHA, significantly inhibited cell viability in a time- and dose-dependent manner^26^. While preclinical trials in patients with primary tumours did not observe a significant effect of MAG-EPA on Ki67 expression in PCa cells, MAG-EPA up-regulated IL-7, which may exert anti-inflammatory effects in the context of PCa^28,29^. These studies suggest that an omega-3-enriched diet has a beneficial effect on primary PCa with potentially no direct consequences on proliferation. Here, supplementing PDOs with EPA showed that EPA alone is insufficient to significantly affect cell viability in PCa compared to normal organoid models. It was also shown that a diet with a high omega-6/omega-3 ratio is associated with an increased risk of high-grade PCa^30^. Accordingly, the recent clinical trial by Aronson and colleagues has shown that increasing the omega-3/omega-6 ratio with fish oil capsules significantly decreased the PCa proliferation index Ki-67^31^. Therefore, increasing the omega-3/omega-6 ratio may help slow PCa progression^31–34^. As such, various studies using distinct models show heterogeneity in their results: EPA sometimes have beneficial effects; sometimes it has no significant effect. Here, we observed that the combination with selenoneine had stronger effects, possibly offering more advantages for patients in upcoming clinical trials.

Several studies have shown that PCa cells can accumulate and use PUFAs from the media to assess different metabolic functions, such as lipogenesis and angiogenesis^29,35,36^. To the best of our knowledge, we are the first to show that both normal prostate and PCa PDO lines can: 1) uptake available fatty acids from the culture media; and 2) process them into downstream intermediates. Notably, treatment with MAG-EPA induces more profound changes in the global lipidomic profile of PDOs. As expected, MAG-EPA, after its uptake by PDOs, was metabolized primarily into omega-3 fatty acids, such as DPA and DHA, in line with increased levels. However, beyond fatty acids, EPA can also be metabolized into different metabolic derivatives. It can be catalyzed by cyclooxygenase-2 (COX-2) and 5-lipoxygenase (LOX-5) into various derivatives, such as prostaglandins and resolvins, to downstream induce anti-inflammatory effects and reduce cell invasion^16,37,38^. How these intermediates are modulated remains to be defined. In addition to generating other omega-3 fatty acids, MAG-EPA increased the omega-3/omega-6 ratio in both normal and tumour PDOs. PDOs metabolize fatty acids from supplemented oils, specifically by synthesizing certain LA isomers in some PCa PDO lines. These isomers may have affinity for the desaturase enzyme and could reduce the conversion of LA into GLA and subsequently AA. These findings indicate that supplementation with omega-3 PUFAs can alter cellular metabolism by decreasing AA concentration, ultimately reducing PCa progression.

Supplementation with MAG-AA resulted in the enrichment of two fatty acids, AA and adrenic acid. Accordingly, adrenic acid metabolism has been shown to induce oxidative stress and produce reactive oxygen species (ROS) in hepatocytes, which are oncogenic in PCa^39^. However, PDOs could metabolize AA into various metabolites different from adrenic acid. Indeed, AA can produce prostaglandins to promote cell growth in PCa. Additionally, 5-LOX can metabolize AA, which may further regulate the *c-Myc* oncogene and facilitate PCa cell proliferation^40,41^. Based on these findings, fatty acids from AA metabolism may promote PCa cell proliferation and potentially lead to more aggressive tumours.

Supplementing PDOs with omega-3 and omega-6 fatty acids produced unique patterns specific to each patient samples, highlighting the high heterogeneity between PDOs (and patients). Such differences may arise from variations in patients’ diets, genetic backgrounds, clinical outcomes, and specific tumour biology. Indeed, both the patients’ genomic profile and the specific genomic alterations of the tumours most probably contribute to such high heterogeneity. In fact, organoids are known to maintain genomic and phenotypic characteristics of Patient-Derived Xenografts (PDXs) from patients with PCa^42^. However, it is essential to understand the optimal exposure time needed for organoids to accurately mimic fatty acid metabolism. Several studies have shown that the response in cell lines to omega-3 is time- and dose-dependent^26,43^. It may be necessary to limit oil supplementation to a maximum of 48 hours for more effective results. Studying the impact of PUFAs through PDOs reveals that patient genomic variability or PDO culture appears to outweigh treatment effects.

In summary, this study is the first to use PDOs as an innovative method for investigating the impact of omega-3 and omega-6 fatty acids on prostate and PCa cell biology. These models, by reconstituting the 3-dimensional structure of the prostate epithelium, offer a valuable new tool added to immortalized cell lines and mouse models. Using these PDOs, we showed that prostate and PCa epithelial cells can take up fatty acids present in the media, metabolize omega-3 fatty acids into downstream intermediates, and that the impact of fatty acid exposure is heterogeneous between patients. Our results using selenoneine also support the hypothesis that combination therapies with MAG-EPA would be more effective than MAG-EPA alone. Future studies are now required to fully appreciate the global impacts of omega-3 or omega-6 nutritional interventions on PCa biology and to fully understand the heterogeneity in the metabolic response of individual tumours.

## Materials and Methods

### Human primary prostate cell culture

The use of human prostate tissues was approved by the Research Ethics Committee of the CHU de Québec – Université Laval (Project 2021-5661). All samples were obtained with the consent of male undergoing radical prostatectomy to treat their prostate cancer.

A series of enzymatic reactions is performed to obtain a single-cell suspension, as described in *Frégeau-Proulx L and al* ^25^. Primary prostate epithelial cells are then seeded in poly-L-lysine-treated culture dishes and cultured in two-dimensional (2D) culture for their amplification in the 2D prostate epithelial cell culture media (Keratinocyte serum-free media (Gibco, 10724-011), Keratinocyte-SFM supplement (Gibco, 37000-015), 10 mM Y-27632 dihydrochloride (Tocris, 1254), 4 ng/mL cholera toxin (Calbiochem Biochemicals, 227035), 500 nM A83-01 (Sigma-Aldrich, SML0788)) ^44^. Cells were incubated at 37°C in 5% CO_2_, and the culture media were changed twice a week. Primary prostate epithelial cells are passaged using TrypLE (Gibco, 12604013) containing 10 mM Y-27632 dihydrochloride, as described in *Jobin C et al* ^44^.

### Human patient-derived organoid culture

Primary prostate epithelial cells are passed as explained previously and resuspended in the 3D prostate epithelial cell culture media (Keratinocyte serum-free media (Gibco, 10724-011), Keratinocyte-SFM supplement (Gibco, 37000-015), 10 mM Y-27632 dihydrochloride (Tocris, 1254), 500 nM A83-01 (Sigma-Aldrich, SML0788)), 10 µM Calcitriol (Cayman Chemical, 71820), 10 nM 17B-estradiol (Sigma-Aldrich, 3301), 10 nM testosterone (Plateforme de chimie médicinale of the CHU de Québec-Universit́é Laval Research Center, Quebec), 2X NeuroCult SM1 neuronal supplement (STEMCELL Technologies, 05711), 1X recombinant mouse Noggin conditioned media (kindly provided by Dr. Alain Veilleux, Université Laval, Quebec), 1X recombinant mouse

R-spondin-1 conditioned media (kindly supplied by Dr. Alain Veilleux, Université Laval, Quebec) ^44^. To obtain a droplet, 75% of Corning® Matrigel® Growth Factor Reduced (GFR) Basement Membrane Matrix, Phenol Red-free, LDEV-free (Corning, 356231) was added to the cells, and 40 µL droplets were seeded into a warm culture plate (24-well plate or 6-well plate, depending on the experiment). Each droplet contained 15000 cells. After 15 min of incubation at 37°C, the 3D prostate epithelial cell culture medium was added to each well. 24 hours after plating, PDOs were treated with selenoneine (1 µg/µL, Pierre Ayotte Lab, Université Laval, Quebec, Canada), HOSO (25 µM, SCF Pharma, Ste-Luce, Canada), MAG-AA (25 µM, NuCheck-Prep Inc, Elysian, MN) and MAG-EPA (25 µM, SCF Pharma). The culture media, with and without treatment, were changed twice a week throughout the experiment, i.e., a 14-day growth period. We used the tumour (T) and normal (N) samples from patients CW488, CW491, CW492, CW513, and CJ71, named PDO#1, PDO#2, PDO#3, PDO#4, and PDO#5.

### Preparation of fatty acid culture media

Stock solutions of MAG-AA and MAG-EPA were prepared by dissolving the fatty acids in DMSO at a concentration of 25 mM. Fatty acids were then dissolved to a final concentration of 25 µM in the 3D prostate epithelial cell culture medium containing 0,25% of BSA (Bio Basic, cat# AD0023). The culture media containing fatty acids were incubated for 60 min at 37°C before being used with PDOs. Tubes were vortexed a few times during the incubation.

### Histology and microscopy analyses

PDO droplets were first fixed in 3% agarose ^25^. PDOs were fixed a second time overnight at room temperature in 10% formalin before being fixed again in paraffin by the *Laboratoire de pathologie de l’Hôtel-Dieu de Québec*. Then, PDOs were cut into 5 mm sections on a microtome (HistoCore MULTICUT 14051856372, Leica). Slides were sent to the pathology department of the Hôtel-Dieu de Québec for Hematoxylin and Eosin (H&E) staining and immunostaining. Images were taken with an EVOS M5000 Imaging System (ThermoFisher Scientific) and the slide scanner ZEISS Axio Scan.Z1 (*Plateforme d’analyses d’images à haut débit du Centre de recherche du CHU de Québec* - Université Laval). Then, organoids were quantified using QuPath v0.6.0^45^.

### Automated PDO quantification

PDO segmentation and quantification were performed on greyscale images using an automated workflow based on the auto-mask feature of the SAM-API extension v0.8^46^ implemented in QuPath software v0.6.0^45^. Organoid annotations generated by the segmentation algorithm were filtered according to greyscale intensity and Haralick sum average thresholds, which were established through validation against manual annotations of a subset of images using receiver operating characteristic (ROC) analysis and area under the curve (AUC) metrics (AUC > 0.8; Youden index). Overlapping annotations were corrected by merging or removing shapes with >50% overlap.

### Cell viability assay

For the cell viability assay, 30 µL droplets containing 75% Matrigel and 11250 cells were seeded into a 96-well black plate. After 14 days of growth and 13 days of treatment, cell viability was assessed using the CellTiter-Glo 3D Cell Viability Assay (Promega, G9681) according to the manufacturer’s protocol. The BioTek Synergy H1 microplate reader was used to read the luminescence.

### Lipidomic / gas chromatography-flame ionization detector / GC-FID

PDOs were cultured for two weeks, both with and without treatment, and then harvested for GC-FID analysis. After harvesting the culture media, the organoids were harvested with an ice-cold 0.9% NaCl solution on ice and then centrifuged for 20 sec at 13,000 g at 4°C. PDOs were washed with an ice-cold 0.9% NaCl solution. Then, the supernatant was discarded, and the pellet was flash-frozen on dry ice and stored at -80 °C. The lipids were extracted from PDOs, as described in various models, such as rats, by Tovar-Parra D and colleagues ^47^. The fatty acid composition of PDOs was determined by gas chromatographic analyses. The pellet was spiked with an internal standard of phosphatidylcholine C:21 (Avanti Polar Lipids, Alabaster, AL). Then, lipids were extracted using a chloroform-methanol mixture (2:1 v/v) according to a modified Folch method^48^. The full fatty acid profiles of lipids were obtained after methylation in methanol/benzene 4:1 (v/v) mixed with acetyl chloride^49^, followed by capillary gas chromatography using a temperature gradient on an HP5890 gas chromatograph (Hewlett Packard, Toronto, Canada). This was equipped with an HP-88 capillary column (100 m x 0.25 mm i.d. x 0.20 µm film thickness; Agilent Technologies) and coupled with a flame ionization detector (FID). Helium was used as a carrier gas (split ratio 1:50). Fatty acids were identified according to their retention times using the following standard mixtures as reference: the FAME 37 mix (Supelco Inc., Bellefonte, PA), the GLC-411 fatty acid mix (NuChek Prep Inc, Elysian, MN), and the methylated fatty acids such as 22:5 ω-6 (Larodan AB, Malmö, Sweden) and 22:5 ω-3 (Supelco Inc., Bellefonte, PA). Additionally, mixtures of trans fatty acids containing cis/trans 18:2 ω-6 (Supelco Inc Bellefonte, PA), and cis/trans 18:3 ω-3 were also incorporated (Supelco Inc Bellefonte, PA)^50^. The lipidomic results were also expressed as a percentage of the total fatty acids.

### Statistical analyses

Statistical analyses were performed in RStudio. Data normality was verified using the Shapiro-Wilk test. Then, a non-parametric test and a post-hoc test were performed using Kruskal-Wallis and Dunnett’s test to compare normal and tumour samples under control conditions, and to compare treatments within normal and tumour conditions. One-way ANOVA followed by Dunnett’s test was performed in GraphPad Prism version 10 to analyze viability data. Heatmap and boxplots were done using complexHeatmap^51^ and ggplot packages.

## Competing Interest Statement

The authors declare no potential conflicts of interest.

## Author contribution

EAW, YF and VF designed the study. CJ and LB performed experiments. MSS did bioinformatic analyses. LL analyzed quantification data in collaboration with GL and KR. PA supplied selenoneine following his patented synthesis process. CM, CA and FP helped obtain prostate tissue for prostatectomy and culturing patient-derived organoids. MSS and EAW analyzed data, made figures and wrote the paper. All authors reviewed the manuscript.

## Acknowledgments

This work was supported by funding to EAW from the Canadian Institutes for Health Research (CIHR; PJT159530). M.S.S. had a scholarship from the Fondation CHU de Québec, the Centre de Recherche sur le Cancer, and the Fonds de recherche du Québec en santé (FRQS). EAW holds a CIHR Tier 2 Canada Research Chair on Metabolic Vulnerabilities of Cancer.

## Notes

### Competing Interest Statement

The authors have declared no competing interest.

## References

1 James, N. D. et al. The Lancet Commission on prostate cancer: planning for the surge in cases. Lancet 403, 1683–1722 (2024). 10.1016/S0140-6736(24)00651-2

2 Down, L. et al. Association between patient ethnicity and prostate cancer diagnosis following a prostate-specific antigen test: a cohort study of 730,000 men in primary care in the UK. BMC Med 22, 82 (2024). 10.1186/s12916-024-03283-5

3 Kimura, T. & Egawa, S. Epidemiology of prostate cancer in Asian countries. Int J Urol 25, 24–531 (2018). 10.1111/iju.13593

4 Shimizu, H. et al. Cancers of the prostate and breast among Japanese and white immigrants in Los Angeles County. Br J Cancer 63, 963–966 (1991). 10.1038/bjc.1991.210

5 Society, C. C. Canadian Cancer Statistics 2025, <https://cancer.ca/en/cancer-information/cancer-types/prostate/statistics> (2025).

6 Kelly, J. et al. Cancer among the circumpolar Inuit, 1989-2003. II. Patterns and trends. Int J Circumpolar Health 67, 408–420 (2008).

7 Friborg, J. T. & Melbye, M. Cancer patterns in Inuit populations. Lancet Oncol 9, 892–900 (2008). 10.1016/S1470-2045(08)70231-6

8 Dewailly, E. et al. n-3 Fatty acids and cardiovascular disease risk factors among the Inuit of Nunavik. Am J Clin Nutr 74, 464–473 (2001). 10.1093/ajcn/74.4.464

9 Zhou, Y. E., Kubow, S. & Egeland, G. M. Highly unsaturated n-3 fatty acids status of Canadian Inuit: International Polar Year Inuit Health Survey, 2007-2008. Int J Circumpolar Health 70, 498–510 (2011). 10.3402/ijch.v70i5.17864

10 Little, M., Achouba, A., Ayotte, P. & Lemire, M. Emerging evidence on selenoneine and its public health relevance in coastal populations: a review and case study of dietary Se among Inuit populations in the Canadian Arctic. Nutr Res Rev 38, 171–180 (2025). 10.1017/S0954422424000039

11 Lemire, M. et al. Local country food sources of methylmercury, selenium and omega-3 fatty acids in Nunavik, Northern Quebec. Sci Total Environ 509-510, 248-259 (2015). 10.1016/j.scitotenv.2014.07.102

12 Achouba, A., Dumas, P., Ouellet, N., Lemire, M. & Ayotte, P. Plasma levels of selenium-containing proteins in Inuit adults from Nunavik. Environ Int 96, 8–15 (2016). 10.1016/j.envint.2016.08.015

13 Shahidi, F. & Ambigaipalan, P. Omega-3 Polyunsaturated Fatty Acids and Their Health Benefits. Annu Rev Food Sci Technol 9, 345–381 (2018). 10.1146/annurev-food-111317-095850

14 Cholewski, M., Tomczykowa, M. & Tomczyk, M. A Comprehensive Review of Chemistry, Sources and Bioavailability of Omega-3 Fatty Acids. Nutrients 10 (2018). 10.3390/nu10111662

15 Innes, J. K. & Calder, P. C. Omega-6 fatty acids and inflammation. Prostaglandins Leukot Essent Fatty Acids 132, 41–48 (2018). 10.1016/j.plefa.2018.03.004

16 Saini, R. K. & Keum, Y. S. Omega-3 and omega-6 polyunsaturated fatty acids: Dietary sources, metabolism, and significance - A review. Life Sci 203, 255–267 (2018). 10.1016/j.lfs.2018.04.049

17 Chevalier, L. & Plourde, M. Comparison of pharmacokinetics of omega-3 fatty acid supplements in monoacylglycerol or ethyl ester in humans: a randomized controlled trial. Eur J Clin Nutr 75, 680–688 (2021). 10.1038/s41430-020-00767-4

18 Liang, P. et al. Effect of omega-3 fatty acid diet on prostate cancer progression and cholesterol efflux in tumor-associated macrophages-dependence on GPR120. Prostate Cancer Prostatic Dis 27, 700–708 (2024). 10.1038/s41391-023-00745-4

19 Gevariya, N. et al. Omega-3 fatty acids decrease prostate cancer progression associated with an anti-tumor immune response in eugonadal and castrated mice. Prostate 79, 9–20 (2019). 10.1002/pros.23706

20 Berquin, I. M. et al. Modulation of prostate cancer genetic risk by omega-3 and omega-6 fatty acids. J Clin Invest 117, 1866–1875 (2007). 10.1172/JCI31494

21 Lachance, G. et al. The gut microbiome-prostate cancer crosstalk is modulated by dietary polyunsaturated long-chain fatty acids. Nat Commun 15, 3431 (2024). 10.1038/s41467-024-45332-w

22 Amaro, G. M. et al. Differential effects of omega-3 PUFAS on tumor progression at early and advanced stages in TRAMP mice. Prostate 82, 1491–1504 (2022). 10.1002/pros.24421

23 Hughes-Fulford, M., Li, C. F., Boonyaratanakornkit, J. & Sayyah, S. Arachidonic acid activates phosphatidylinositol 3-kinase signaling and induces gene expression in prostate cancer. Cancer Res 66, 1427–1433 (2006). 10.1158/0008-5472.CAN-05-0914

24 Aronson, W. J. et al. Phase II prospective randomized trial of a low-fat diet with fish oil supplementation in men undergoing radical prostatectomy. Cancer Prev Res (Phila) 4, 2062–2071 (2011). 10.1158/1940-6207.CAPR-11-0298

25 Frégeau-Proulx, L. et al. Multiple metabolic pathways fuel the truncated tricarboxylic acid cycle of the prostate to sustain constant citrate production and secretion. Mol Metab 62, 101516 (2022). 10.1016/j.molmet.2022.101516

26 Sun, Y., Jia, X., Hou, L., Liu, X. & Gao, Q. Involvement of apoptotic pathways in docosahexaenoic acid-induced benefit in prostate cancer: Pathway-focused gene expression analysis using RT. Lipids Health Dis 16, 59 (2017). 10.1186/s12944-017-0442-5

27 Yu, K. J. et al. EPA Modulates KLK Genes via miR-378: A Potential Therapy in Prostate Cancer. Cancers (Basel) 14 (2022). 10.3390/cancers14112813

28 Robitaille, K. et al. A phase IIb randomized placebo-controlled trial testing the effect of MAG-EPA long-chain omega-3 fatty acid dietary supplement on prostate cancer proliferation. Commun Med (Lond) 4, 56 (2024). 10.1038/s43856-024-00456-4

29 Gevariya, N. et al. Omega-3 Eicosapentaenoic Acid Reduces Prostate Tumor Vascularity. Mol Cancer Res 19, 516–527 (2021). 10.1158/1541-7786.MCR-20-0316

30 Williams, C. D. et al. A high ratio of dietary n-6/n-3 polyunsaturated fatty acids is associated with increased risk of prostate cancer. Nutr Res 31, 1–8 (2011). 10.1016/j.nutres.2011.01.002

31 Aronson, W. J. et al. High Omega-3, Low Omega-6 Diet With Fish Oil for Men With Prostate Cancer on Active Surveillance: The CAPFISH-3 Randomized Clinical Trial. J Clin Oncol 43, 800–809 (2025). 10.1200/JCO.24.00608

32 Aronson, W. J. et al. Modulation of omega-3/omega-6 polyunsaturated ratios with dietary fish oils in men with prostate cancer. Urology 58, 283–288 (2001). 10.1016/s0090-4295(01)01116-5

33 Apte, S. A., Cavazos, D. A., Whelan, K. A. & Degraffenried, L. A. A low dietary ratio of omega-6 to omega-3 Fatty acids may delay progression of prostate cancer. Nutr Cancer 65, 556–562 (2013). 10.1080/01635581.2013.775316

34 Kobayashi, N. et al. Effect of altering dietary omega-6/omega-3 fatty acid ratios on prostate cancer membrane composition, cyclooxygenase-2, and prostaglandin E2. Clin Cancer Res 12, 4662–4670 (2006). 10.1158/1078-0432.CCR-06-0459

35 Balaban, S. et al. Extracellular Fatty Acids Are the Major Contributor to Lipid Synthesis in Prostate Cancer. Mol Cancer Res 17, 949–962 (2019). 10.1158/1541-7786.MCR-18-0347

36 Watt, M. J. et al. Suppressing fatty acid uptake has therapeutic effects in preclinical models of prostate cancer. Sci Transl Med 11 (2019). 10.1126/scitranslmed.aau5758

37 Hyde, C. A. & Missailidis, S. Inhibition of arachidonic acid metabolism and its implication on cell proliferation and tumour-angiogenesis. Int Immunopharmacol 9, 701–715 (2009). 10.1016/j.intimp.2009.02.003

38 Panagiotopoulos, A. A., Kalyvianaki, K., Castanas, E. & Kampa, M. Eicosanoids in prostate cancer. Cancer Metastasis Rev 37, 237–243 (2018). 10.1007/s10555-018-9750-0

39 Zhao, J. et al. Adrenic acid induces oxidative stress in hepatocytes. Biochem Biophys Res Commun 532, 620–625 (2020). 10.1016/j.bbrc.2020.08.102

40 Monga, J., Subramani, D., Bharathan, A. & Ghosh, J. Pharmacological and genetic targeting of 5-lipoxygenase interrupts c-Myc oncogenic signaling and kills enzalutamide-resistant prostate cancer cells via apoptosis. Sci Rep 10, 6649 (2020). 10.1038/s41598-020-62845-8

41 Ghosh, J. & Myers, C. E. Arachidonic acid stimulates prostate cancer cell growth: critical role of 5-lipoxygenase. Biochem Biophys Res Commun 235, 418–423 (1997). 10.1006/bbrc.1997.6799

42 Beshiri, M. L. et al. A PDX/Organoid Biobank of Advanced Prostate Cancers Captures Genomic and Phenotypic Heterogeneity for Disease Modeling and Therapeutic Screening. Clin Cancer Res 24, 4332–4345 (2018). 10.1158/1078-0432.CCR-18-0409

43 LeMay-Nedjelski, L., Mason-Ennis, J. K., Taibi, A., Comelli, E. M. & Thompson, L. U. Omega-3 Polyunsaturated Fatty Acids Time-Dependently Reduce Cell Viability and Oncogenic MicroRNA-21 Expression in Estrogen Receptor-Positive Breast Cancer Cells (MCF-7). Int J Mol Sci 19 (2018). 10.3390/ijms19010244

44 Jobin, C. et al. Protocol for transducing human primary epithelial prostate cells and patient-derived organoids with high efficiency. STAR Protoc 5, 103200 (2024). 10.1016/j.xpro.2024.103200

45 Bankhead, P. et al. QuPath: Open source software for digital pathology image analysis. Sci Rep 7, 16878 (2017). 10.1038/s41598-017-17204-5

46 Ko, S. Training deep learning models for cell image segmentation with sparse annotations. bioRxiv (2023). 10.1101/2023.06.13.544786

47 Tovar-Parra, D. et al. The rat mammary gland undergoes dynamic transcriptomic and lipidomic modifications from pre-puberty to adulthood. Sci Rep 15, 12222 (2025). 10.1038/s41598-025-97532-z

48 Shaikh, N. A. & Downar, E. Time course of changes in porcine myocardial phospholipid levels during ischemia. A reassessment of the lysolipid hypothesis. Circ Res 49, 316–325 (1981). 10.1161/01.res.49.2.316

49 Lepage, G. & Roy, C. C. Direct transesterification of all classes of lipids in a one-step reaction. J Lipid Res 27, 114–120 (1986).

50 Simard, M. et al. Postmortem Fatty Acid Abnormalities in the Cerebellum of Patients with Essential Tremor. Cerebellum 23, 2341–2359 (2024). 10.1007/s12311-024-01736-4

51 Gu, Z., Eils, R. & Schlesner, M. Complex heatmaps reveal patterns and correlations in multidimensional genomic data. Bioinformatics 32, 2847–2849 (2016). 10.1093/bioinformatics/btw313

